# ASO targeting temperature-controlled *RBM3* poison exon splicing prevents neurodegeneration in vivo

**DOI:** 10.1101/2022.10.26.513170

**Authors:** Marco Preußner, Heather L Smith, Min Zhang, Daniel Hughes, Ann-Kathrin Emmerichs, Silvia Scalzitti, Diego Peretti, Dean Swinden, Alexander Neumann, Tom Haltenhof, Giovanna R Mallucci, Florian Heyd

## Abstract

Neurodegenerative diseases are increasingly prevalent in the aging population, yet currently no disease-modifying treatments are available. Increasing the expression of the cold-shock protein, RBM3, through therapeutic hypothermia is remarkably neuroprotective, but cooling poses a health risk itself, strongly limiting its clinical application. Selective upregulation of RBM3 at normothermia thus holds immense therapeutic potential. Here we identify a poison exon within the RBM3 gene that is solely responsible for cold-induced RBM3 expression. Genetic removal or ASO-mediated manipulation of this exon yields high RBM3 levels independent of cooling. Notably, a single administration of ASO to exclude the poison exon, using FDA-approved chemistry, results in long-lasting increase of RBM3 expression in mouse brains. In prion-diseased mice, this treatment leads to remarkable neuroprotection, with prevention of neuronal loss and spongiosis despite high levels of prion protein. RBM3-inducing ASOs could thus broadly deliver protection in humans in conditions ranging from acute brain injury to Alzheimer’s disease.

**One sentence summary:** Inducing cold shock protein RBM3 by modulating its alternative splicing at normothermia is neuroprotective in vivo

## Main

The expression of glycine-rich RNA binding proteins upon cooling, such as CIRBP (Cold-induced RNA binding protein) and RBM3 (RNA binding motif-3), was originally described in the 1990s (Danno *et al*, 1997; Nishiyama *et al*, 1997). However, despite evolutionary conservation (Ciuzan *et al*, 2015) and extreme temperature sensitivity of this phenomenon (Jackson *et al*, 2015; Los *et al*, 2022), the mechanistic basis for cold-induced RBM3 expression has remained enigmatic. Our recent identification of temperature-regulated alternative splicing coupled to nonsense-mediated decay (NMD) provides a global mechanism for the control of temperature-dependent gene expression (Neumann *et al*, 2020). The NMD pathway recognizes mRNA isoforms containing premature termination codons (PTCs) and targets these mRNAs for degradation, thus allowing splicing-controlled regulation of gene expression (Lykke-Andersen & Jensen, 2015). NMD-inducing poison isoforms are frequently found in RNA binding proteins (Neumann *et al*., 2020) and exclusion of a poison exon upon cooling in CIRBP provides an explanation for cold-induced expression (Haltenhof *et al*, 2020). While CIRBP has diverse functions ranging from circadian sleep homeostasis to inflammation and cancer (Hoekstra *et al*, 2019; Lujan *et al*, 2018; Morf *et al*, 2012; Qiang *et al*, 2013), the closely related RBM3 protein is strongly associated with the neuroprotective effect of hypothermia. This has been observed in scenarios ranging from *in vitro* protection from forced apoptosis of neuronal cell lines and brain slices (Chip *et al*, 2011) to profound neuroprotective effects *in vivo*, where RBM3 induction by cooling or over-expression restores memory, prevents synapse and neuronal loss and extends survival in preclinical mouse models of prion and Alzheimer’s disease (Peretti *et al*, 2015; Peretti *et al*, 2021). More recently, RBM3 has been shown to stimulate neurogenesis in rodent brain after hypoxic-ischemic brain injury (Zhu *et al*, 2019) and to protect against neurotoxin effects in neuronal cell lines (Yang *et al*, 2019). Therapeutic hypothermia is used in some clinical settings for neuroprotection, including neonatal hypoxic ischemic encephalopathy, head injury (Azzopardi *et al*, 2014; Shankaran, 2012; Shankaran *et al*, 2005; Thayyil *et al*, 2021), stroke and during cardiac surgery in adults (2002; Bernard *et al*, 2016; Dankiewicz *et al*, 2021; Lascarrou *et al*, 2019; Nielsen *et al*, 2013). While the mechanisms of hypothermia-induced neuroprotection in humans are not fully understood, RBM3 is used as a biomarker of success in intensive care unit (ICU) patients exposed to hypothermia (Rosenthal *et al*, 2019). Induced cooling in humans requires an ICU set up and is not without risk, with high prevalence of blood clots, pneumonia, and other complications, strongly limiting its clinical use. Inducing RBM3 without cooling would bypass these risks and requirements and could represent a much needed new neuroprotective strategy broadly applicable in conditions ranging from stroke and brain injury to Alzheimer’s and other neurodegenerative diseases. Therefore, a detailed mechanistic understanding of cold-induced RBM3 expression offers immense therapeutic potential.

We hypothesized that cold-induced RBM3 expression, similar to CIRBP expression (Haltenhof *et al*., 2020), could be regulated via cold-induced exclusion of a poison isoform that is present at normal or high temperatures. While a global splicing analysis in primary mouse hepatocytes did not reveal temperature-controlled alternative splicing in *rbm3* (Neumann *et al*., 2020), a more focused analysis revealed an uncharacterized exon containing seven PTCs within the evolutionarily conserved intron 3. This exon, which we call exon 3a (E3a), is not included in the mRNA at colder temperature (34°C), but E3a containing isoforms are detectable at warm temperature (38°C) and become strongly stabilized upon NMD inhibition via cycloheximide (CHX) (Figure 1A). Using splicing sensitive radioactive RT-PCRs we confirmed a very similar temperature-controlled E3a splicing pattern in mouse primary hippocampal neurons (Figure 1B). Consistent with high evolutionary conservation (Figure 1A, bottom) and an almost identical E3a sequence in humans we also find temperature-controlled inclusion of the human *RBM3* E3a homolog. In HEK293 cells E3a inclusion is highly temperature responsive within the physiologically relevant temperature range between 33°C to 39°C, correlating with highly temperature-sensitive *RBM3* expression (Figures 1C, 1D and S1A). In addition, *RBM3* E3a inclusion responds quickly, within 4 hours, to an external square wave temperature rhythm (Figure S1B) and could therefore control *RBM3* expression in response to circadian body temperature changes (Liu *et al*, 2013; Morf *et al*., 2012). The presence of seven PTCs within E3a (Figure 1A) and the strong stabilization upon addition of the translation inhibitor CHX suggest NMD-mediated degradation of the E3a isoform. To provide further evidence for degradation via the NMD pathway, we investigated stabilization of the E3a-containing isoform in response to NMD factor knockdowns (Colombo *et al*, 2017). In human cells, the *RBM3* E3a isoform is reversibly stabilized upon *UPF1* knockdown and dramatically and reversibly stabilized (∼75% NMD isoform) upon *SMG6/7* double knockdown (Figure 1E and S1C). In summary, these data identify an uncharacterized poison exon in *RBM3* that could control cold-induced *RBM3* expression through cooling induced skipping and evasion of NMD.

**Figure 1:**
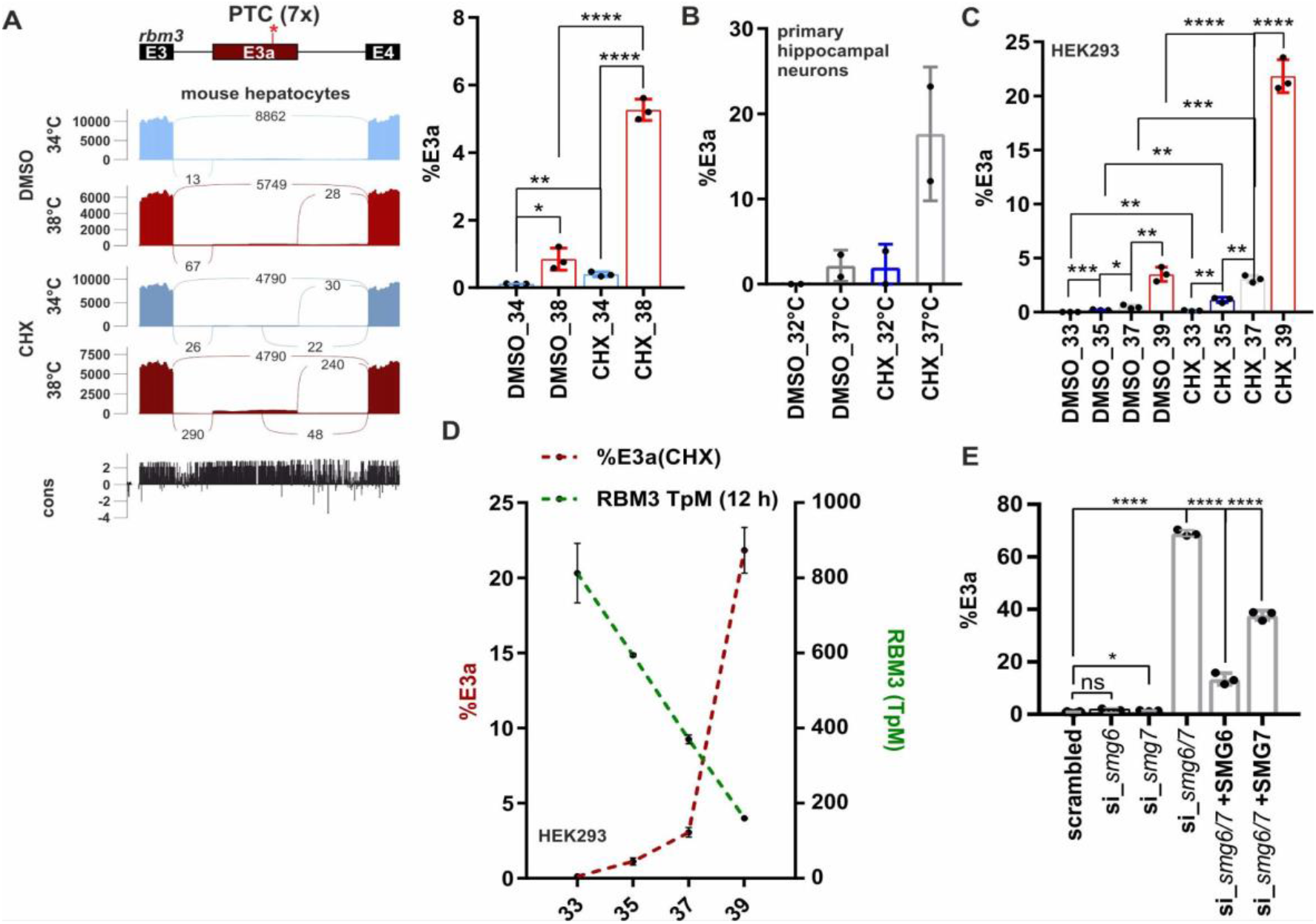
*RBM3* intron 3 contains an evolutionary conserved heat-induced poison exon. **A**. Sashimi blot identifies an uncharacterized exon (E3a; with 7 premature termination codons: PTC) within *rbm3* intron 3. Mouse primary hepatocytes were incubated at 34°C or 38°C with or without the translation inhibitor cycloheximide (CHX, DMSO as solvent control) and analysed by RNA sequencing. Below the simplified exon-intron structure, the Sashimi plot shows the distribution of raw sequencing reads. Exon-Exon junction reads are indicated by the numbers connecting the exons. At the bottom, high sequence conservation across placental species is indicated. Quantification of %E3a inclusion in RNA sequencing samples is shown on the right (mean ± SD, n=3, all individual data points are shown). **B**. Quantification of radioactive splicing sensitive RT-PCRs confirm heat-induced and CHX-stabilized formation of the E3a containing isoform at warmer temperatures in primary hippocampal neurons (mean ± SD, n=2, all individual data points are shown). **C**. *RBM3* E3a regulation is conserved in humans. HEK293 cells were incubated at the indicated temperatures for 12h (DMSO/CHX last 4h) and investigated for E3a inclusion as in B (mean ± SD, n=3, all individual data points are shown). For a representative gel image see also Figure S1A. **D**. Gene expression of *RBM3* anti-correlates with inclusion of E3a. Transcripts per million (TpM) values for *RBM3* are derived from RNA sequencing data from HEK293 cells incubated for the indicated time points for 12 hours and are plotted on the right y-axis (green, n=2, mean ± SD). Inclusion levels for E3a are derived from C. **E**. *RBM3* E3a stabilization in response to *SMG6* and *SMG7* knockdown and rescue (mean ± SD, n=3, all individual data points are shown). In all panels, statistical significance was determined by unpaired t-test and is indicated by asterisks: p values: *p<0.05, **p<0.01, ***p<0.001, ****p<0.0001. Data from (Colombo *et al*., 2017).

To address an involvement of E3a in temperature-controlled *RBM3* expression, we used CRISPR/Cas9-mediated genome editing to generate cell lines lacking *RBM3* E3a. After clonal selection we obtained two homozygous cell lines, derived from distinct guide RNA pairs (Figures 2A and S2A). While two control cell lines, generated from px459 empty vector transfected cells, showed strong temperature controlled *RBM3* mRNA and protein expression, this was abrogated in cell lines lacking E3a (Figures 2B, 2C and S2B). Importantly, cell lines lacking E3a did not only lose *RBM3* temperature sensitivity but showed a constantly high *RBM3* expression level, that in control cells was only reached at low temperature. These data suggest that temperature-controlled alternative splicing coupled to NMD is the main mechanism that controls *RBM3* expression levels in the physiologically relevant temperature range. Consistent with this, recent work has implicated several splicing factors, amongst others HNRNPH1, in regulating *RBM3* expression through poison exon splicing (Lin *et al*, 2022). Temperature-dependent phosphorylation of SR proteins likely contributes to this regulation (Haltenhof *et al*., 2020; Preussner *et al*, 2017), as the effect of temperature on *RBM3* expression is strongly reduced in conditions with inhibited CDC-like kinases (CLKs; Figure S2C). Importantly, this mechanism offers the possibility to manipulate RBM3 expression by modulating alternative splicing of E3a using antisense oligonucleotides (ASOs). A splice-modulating ASO is already in clinical use to treat spinal muscular atrophy (SMA) in humans (Finkel *et al*, 2016) thus representing an established approach to therapeutically manipulate alternative splicing in the human nervous system with beneficial effect. As proof of principle, we used a splice-site blocking morpholino (MO) directed against the 5’ss of *RBM3* E3a. This MO induced *RBM3* mRNA levels substantially in HEK293 cells, confirming that *RBM3* expression can be controlled in *trans* by targeting splicing of E3a (Figure 2D; see also below). Importantly, MO transfection also induced RBM3 protein expression in mouse primary hippocampal neurons at 37°C (Figures 2E and S2D), validating our findings in a setup relevant for neuroprotection/degeneration. Together, these data provide strong evidence for splicing-controlled *RBM3* expression and identify E3a as a potential therapeutic target for a neuroprotective increase of RBM3 expression at normothermia.

**Figure 2:**
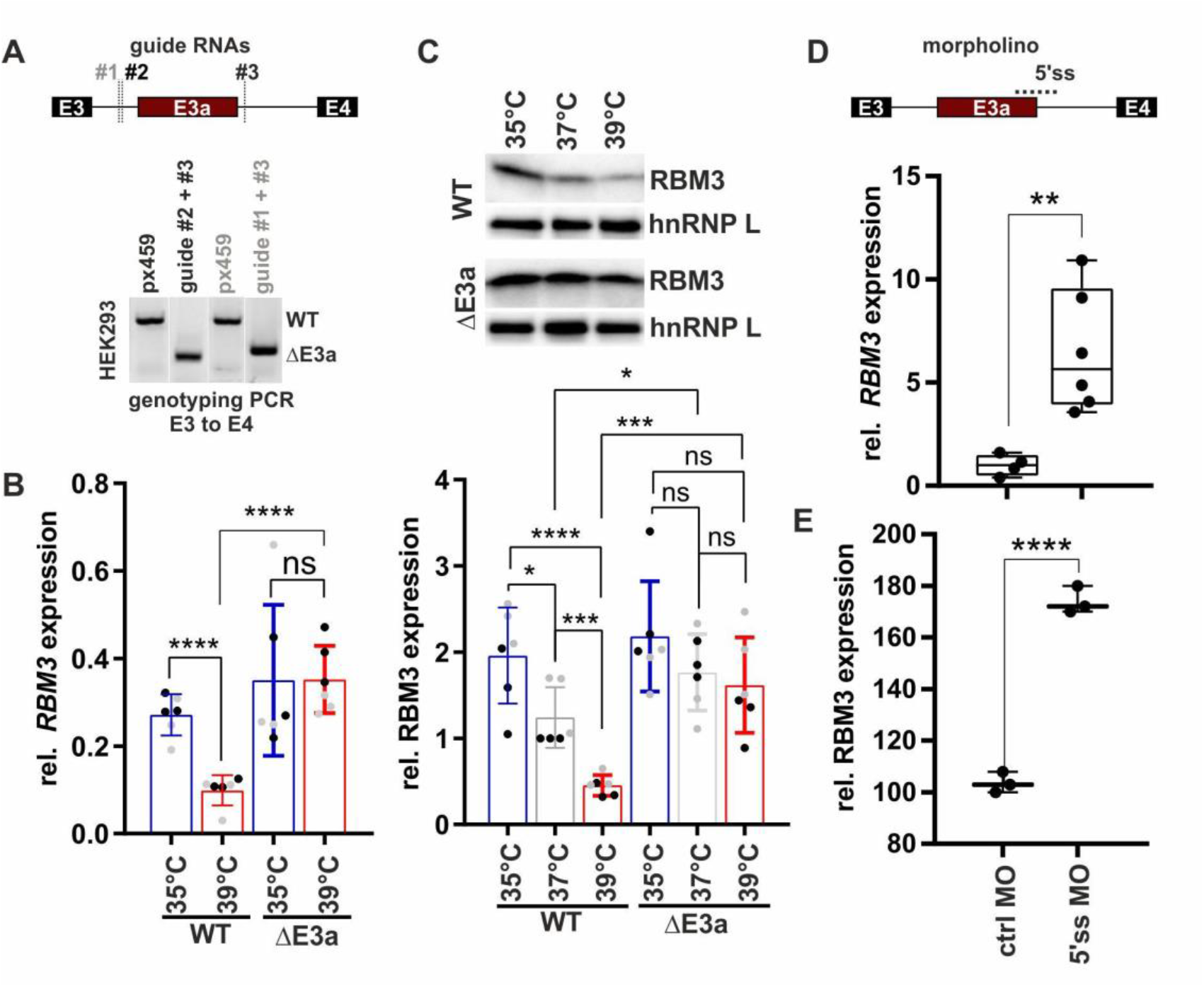
E3a controls temperature dependent RBM3 expression. **A**. CRISPR/CAS9 mediated removal of *RBM3* E3a. One of two guide RNAs targeting the upstream intron (#1, #2) were co-transfected with a guide RNA targeting the downstream intron (#3). Below, genotyping PCR after clonal selection with px459 transfected cells serving as a negative control. See also Figure S2A. **B, C**. RT-qPCR (B) and Western blot (C) analysis of *RBM3* levels in edited cell lines. Clonal cell lines from A were incubated at the indicated temperatures for 24h. In B, isolated RNA was investigated by qPCR and *rbm3* expression is shown relative to *GAPDH* levels. In C, lysates from an independent experiment were investigated for RBM3 protein expression, hnRNP L served as a loading control, a representative gel (left) and quantification (right) are shown (mean ± SD, n=6 (3 per clone, indicated in black/grey), all individual data points are shown). Statistical significance was determined by unpaired t-tests and is indicated by asterisks *p<0.05, ***p<0.001, ****p<0.0001. See Figure S2B for all gels. **D, E**. Manipulation of *RBM3* E3a splicing directly controls *RBM3* expression levels. In D, HEK293 cells were transfected for 48h at 37°C with a MO blocking the 5’ss of E3a. *RBM3* expression is shown relative to *GAPDH* levels and normalized to a non-targeting MO. Statistical significance was determined by unpaired t-test and is indicated by asterisks: p values: **p<0.01 (n=4-6). In E, primary hippocampal neurons were transfected with the indicated MOs for 48 hours and investigated for RBM3 by Western blotting. GAPDH served as a loading control (n=3; unpaired t-test derived p-value ****p<0.0001). Gel images in Figure S2D.

Manipulation of (non-productive) alternative splicing with ASOs has broad therapeutic potential (Bennett *et al*, 2019; Lim *et al*, 2020), but the efficiency of induced exon skipping is strongly dependent on the exact sequence and chemistry of the oligonucleotide (Erdos *et al*, 2021; Puttaraju *et al*, 2021). Therefore, the design of the most potent ASOs often requires intensive screening. To narrow down potential target sites for antisense-based therapeutics, we started with a minigene analysis. In a minigene context, NMD isoforms are not degraded as the minigene-derived RNA is not translated, and minigenes allow systematic mutagenesis to decipher *cis*-regulatory elements. We cloned the human or mouse genomic sequence comprising *RBM3* exons 3 to 4 and analysed minigene splicing after transfection of (human) HEK293 (Figures 3A, 3B and S3A) and (mouse) neuroblastoma N2a cells, respectively (Figure S3A). Similar to endogenous splicing, minigene splicing responds to temperature in a gradual manner (Figure 3B) and is indistinguishable between mouse and human minigenes (Figure S3A), indicating that the minigene contains all *cis*-regulatory elements required for evolutionarily conserved temperature-controlled alternative splicing. Screening mutagenesis by replacing 50 nucleotide windows with human *beta*-globin sequences revealed two strong enhancer elements, both in HEK293 at 37 or 39°C (M2 and M4; Figures 3C and S3B, Table S1). We also identified silencer elements in the regions downstream of the 3’ss (M1), close to an internal 3’ss (M3) and upstream of the 5’ss (M6). In contrast to mutations of the M2 and M4 enhancer elements or deleting their evolutionary conserved core sequence, silencer mutants remained mostly temperature sensitive (compare 37°C and 39°C in Figure S3B, and S3C). This suggests that temperature sensitivity is mainly mediated by the enhancers, whereas the silencer elements rather control basal E3a inclusion levels, for example by controlling splice site accessibility. Deleting (instead of replacing) the M2 or M4 enhancer sequences also abolished E3a inclusion, and smaller replacements revealed the location of the enhancer elements within nucleotides 42 to 71 and 142 to 171 of the exon (Figure S3D and Table S1).

**Figure 3:**
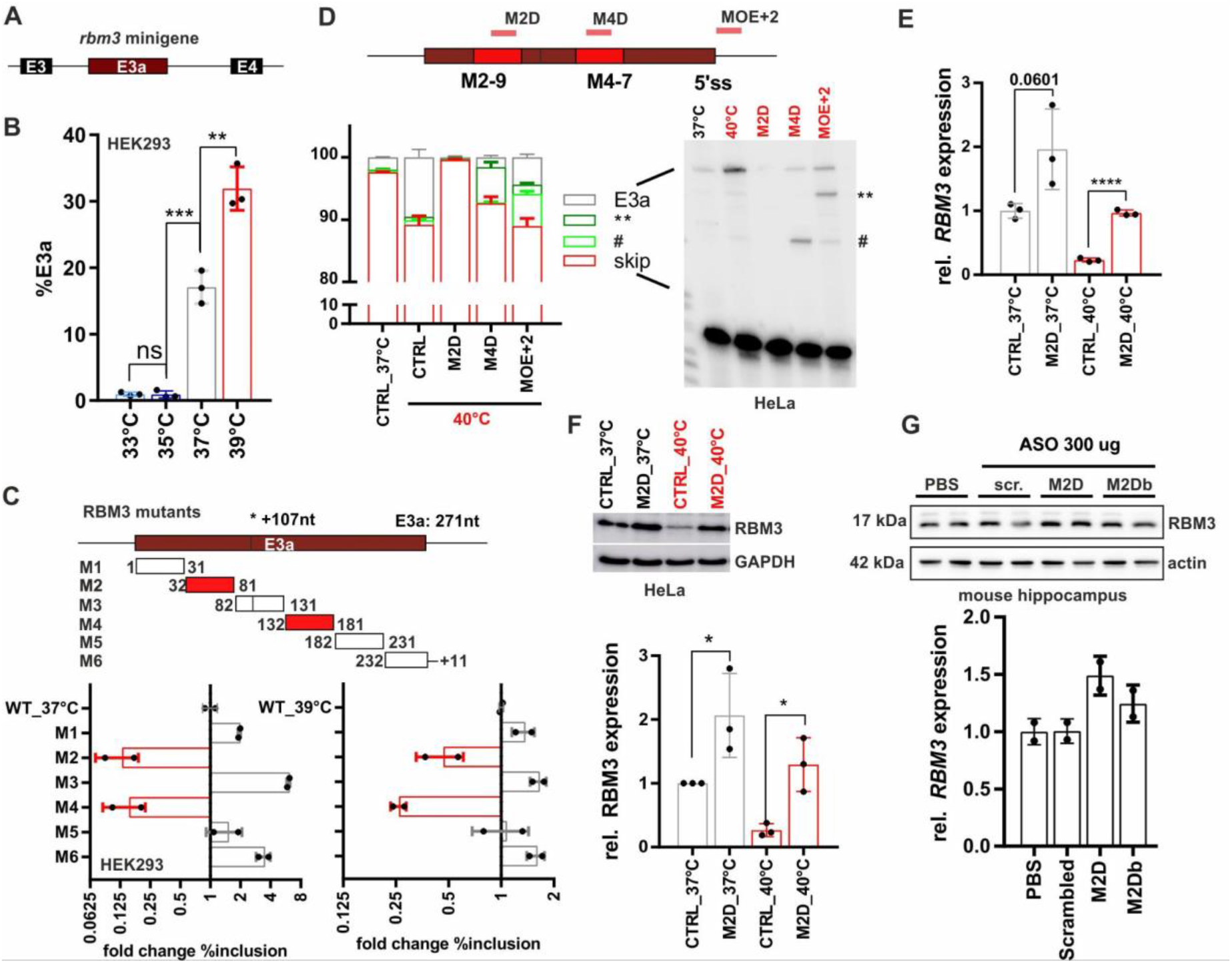
Targeting enhancer elements to control *RBM3* E3a inclusion. **A, B**. An *rbm3* minigene reproduces temperature controlled E3a inclusion. In A the minigene structure containing the whole unshorten sequence from E3 to E4 (including upstream 3’ss and downstream 5’ss) is shown. In (B) minigenes from mouse were transfected into HEK293 and incubated at the indicated temperatures for 12h. E3a inclusion was investigated by splicing sensitive PCR and quantified, %E3a is shown (mean ± SD, n=3, all individual data points are shown; unpaired t-test derived p-value **p<0.01, ***p<0.001). **C**. Systematic mutational screening for regulatory elements. Mutations resulting in exon skipping are highlighted in red. Below, analysis of fold change in %E3a in HEK293 at 37°C and 39°C (mean ± SD, n=2). Position 107 marks an alternative 3’ss in E3a. See also Figure S3B and Table S1. **D**. ASOs targeting M2-9, M4-7 or the 5’ss (see Figure S4A and Table S2) prevent endogenous *RBM3* E3a inclusion in human HeLa cells. ASO-transfected cells were kept for 24 hours at 40°C. Control samples at 37°C and 40°C are shown; CHX was added for the last 4 hours. Exon 3a inclusion was investigated by splicing sensitive RT-PCR, a representative gel and phosphorimager quantification are shown (mean ± SD, n=3). The hashtag marks the use of internal 5’ and 3’ss that is promoted by all ASOs targeting the M4 region. ASOs targeting the 5’ss induced the usage of an internal 5’ss (marked by two asterisks). **E, F**. M2D induces RBM3 mRNA (E) and protein (F) expression in human HeLa cells. ASOs were transfected for 24 hours at 37°C (grey) or at 40°C (red). RBM3 induction was measured relative to GAPDH expression (mean ± SD, n=3, all individual data points are shown; unpaired t-test derived p-value **p<0.01, ****p<0.0001). **G**. M2D induces RBM3 protein expression *in vivo*. Hippocampus samples from two independent mice per condition were analyzed by Western blotting (left) and RBM3 protein was quantified relative to actin and PBS (right, n=2).

As a basis for therapeutic manipulation of RBM3 levels in humans, we used this minigene analysis and performed an ASO screen targeting different cis-regulatory elements. We identified several ASOs targeting the M2 or M4 enhancers or the 5’ss that prevent E3a inclusion in HeLa cells (Figures 3D and S4A-C, Table S2). We noticed that ASOs targeting the M4 region or the 5’ss result in partial usage of internal alternative 3’ or 5’ splice sites leading to products that could still induce NMD. However, all variants targeting the M2D region quantitatively abolish E3a inclusion (Figure S4C), making them promising candidates for therapeutic applications. Importantly, the M2D ASO induces RBM3 expression at 40°C to the level observed in control cells at 37°C, and is also inducing RBM3 expression up to 2 fold at 37°C (Figures 3E and 3F). M2D and M2Db also worked in a mouse cell line, as they induced *Rbm3* expression 1.5 (37°C) to 4 (39°C) fold in N2a cells (Figure S4D). In these experiments, we combined a phosphorothioate (PS) modified backbone with uniform 2′-O-methoxyethyl (MOE) modified bases as in the FDA approved drug Nusinersen (Hua *et al*, 2011), which allows distribution throughout the central nervous system after intrathecal injection (Finkel *et al*., 2016) and systemic delivery in vivo (Sheng *et al*, 2020). To provide *in vivo* evidence that AS-NMD modulating ASOs can increase RBM3 levels in a therapeutic range in the central nervous system, we chose two ASOs targeting the M2 region, M2D and M2Db. We first checked for efficacy in RBM3 induction of the individual ASOs when administered by a single intracerebroventricular injection to wild type mice at doses of 100μg and 300μg. Both ASOs were well tolerated up to 3 weeks post injection and both increased RBM3 protein levels. M2D was the more efficient, resulting in 1.5-fold increase of RBM3 in the hippocampus at both 100μg and 300μg doses (Figures 3G and S4E) and correlating with reduced E3a inclusion levels (Figure S4F). We therefore focused on M2D for further testing *in vivo*.

We tested the therapeutic potential of M2D-mediated RBM3 induction in a mouse prion disease model extensively used to test the effects of cooling and RBM3 over-expression on the progression of neurodegeneration (Bastide *et al*, 2017; Peretti *et al*., 2015; Peretti *et al*., 2021). Hemizygous tg37+/− mice overexpress prion protein (PrP) at around 3-fold over wild-type levels (Mallucci *et al*, 2002). When inoculated with Rocky Mountain Laboratory (RML) prions, these mice show a rapid incubation time, succumbing to disease in only 12 weeks post inoculation (w.p.i.), with rapidly progressing spongiform change and extensive neurodegeneration throughout the brain, including hippocampal CA1-3 regions (Mallucci *et al*., 2002). Early cooling (at 3 w.p.i.) to boost RBM3 levels, or lentiviral delivery of RBM3 to the hippocampus, are both profoundly neuroprotective in prion-diseased tg37 mice and in Alzheimer’s 5xFAD mice, whereas RNAi of RBM3 eliminates the protective effects of cooling in both models (Peretti *et al*., 2015). We treated prion-diseased tg37 mice (n=8) with 200 μg of the M2D ASO or a scrambled ASO control (Figure 4A). ASOs were delivered by a single intracerebroventricular injection at 3 w.p.i., consistent with the timing of our previous interventions (Bastide *et al*., 2017; Peretti *et al*., 2015; Peretti *et al*., 2021). Mice treated with M2D/scrambled ASO were analyzed for neuroprotection at 12 w.p.i. when all scrambled-ASO treated mice had succumbed to prion disease. Remarkably, the single dose of M2D ASO resulted in RBM3 levels 2-fold higher than scrambled-treated mice 9 weeks after injection, at 12 w.p.i. (Figure 4B). Higher levels of RBM3 were associated with marked neuroprotection: 7/8 M2D-treated mice showed extensive preservation of pyramidal neurons in the hippocampal CA1-3 regions (Figure 4C and 4D) compared to scrambled-ASO treated mice, of which 5 of 6 mice showed profound neuronal loss in these regions (Figures 4C and 4D). M2D-mediated neuroprotection was associated with markedly lower spongiosis scores compared to scrambled-treated mice (Figure 4E) and did not alter total levels of PrP or of the amyloid form, proteinase K-resistant PrP^Sc^ (Figure 4F). These data support a robust and long-lasting induction of RBM3 without cooling by a single dose of M2D, with remarkable neuroprotection in the context of a rapidly progressive neurodegenerative disorder. The ability to raise RBM3 levels by one administration of a well-tolerated ASO using FDA-approved chemistry, in place of therapeutic hypothermia, has strong implications for neuroprotection in diverse conditions, from acute treatment of neonates through to cardiac surgery, stroke and head injury in adults, to longer term neuroprotection in degenerative disorders. In the acute setting, this approach yields the neuroprotective effect of RBM3 while bypassing the substantial risks associated with intensive care and hypothermia. Further, the approach has marked appeal in the prevention of a variety of neurodegenerative disorders. ASOs are highly successful in children with SMA, and have been recently licensed for the treatment of the rapidly progressive adult neurodegenerative disorder, ALS (Miller *et al*, 2022). In the search for disease-modifying therapies for Alzheimer’s disease and related dementias, induction of the broadly neuroprotective RBM3 via ASO delivery to drive its long-term expression is a compelling therapeutic approach. Increasing RBM3 expression boosts neuronal resilience and synapse regeneration that are key to resisting the direct and indirect toxic effects of protein misfolding disorders, particularly in the context of age-related dementias.

**Figure 4:**
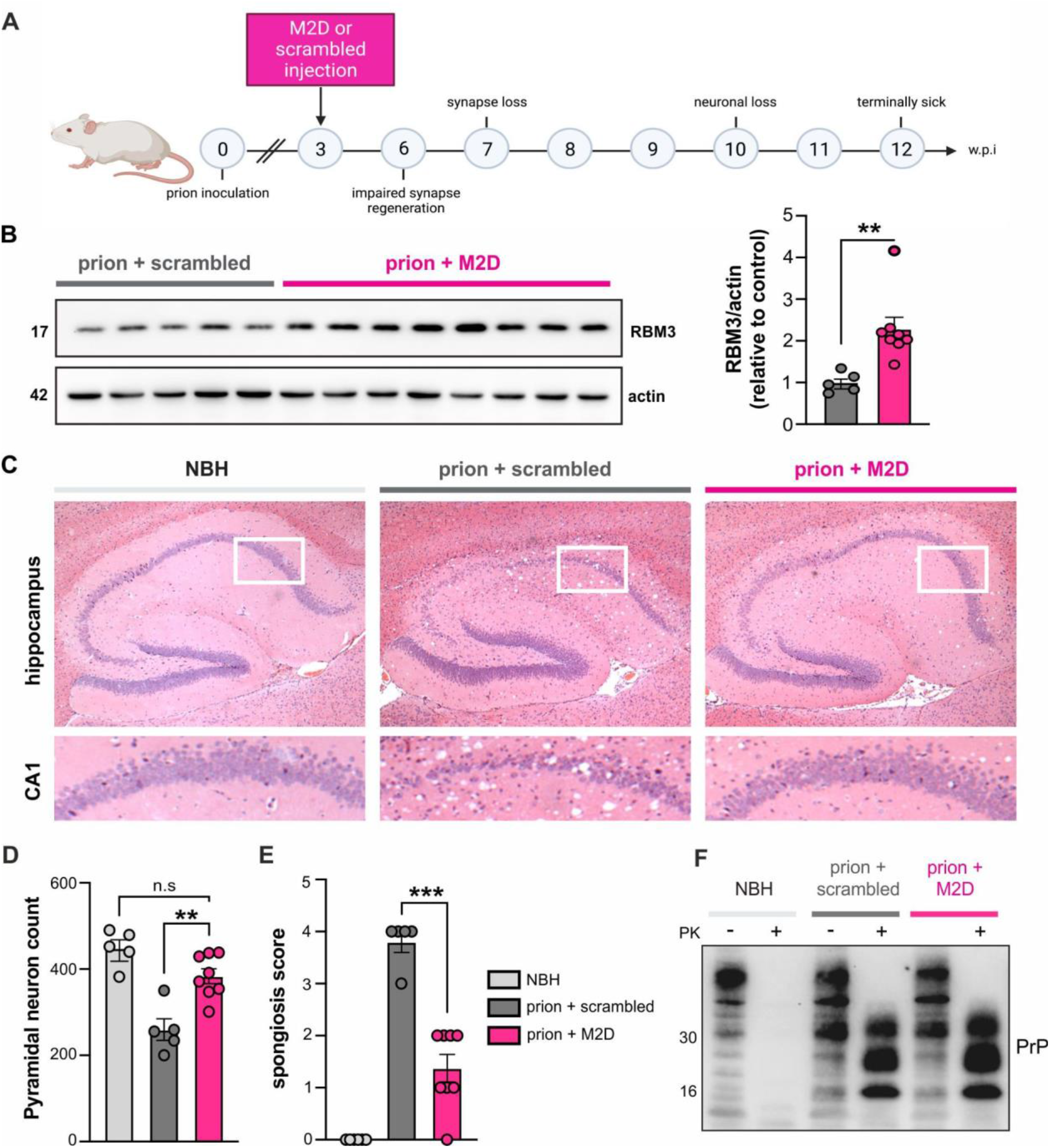
M2D elevates hippocampal RBM3 levels and is profoundly neuroprotective. **A**. Schematic of experimental design. Prion-inoculated tg37+/− mice were injected with 200 μg of either M2D or a scrambled ASO at 3 w.p.i. **B**. Western blot of hippocampal lysates from prion-infected mice treated with scrambled ASO or M2D. M2D increases RBM3 expression by 2-fold compared to scrambled-treated mice, 9 weeks after ASO injection at 12 w.p.i. **C**. Representative images of haematoxylin and eosin stained brain slices from NBH (control), and prion-infected scrambled- and M2D-treated mice at 12 w.p.i. when scrambled-treated mice were culled for prion signs. M2D confers marked neuroprotection in the hippocampus, with conservation of CA1-3 pyramidal layer, protection from shrinkage of the whole hippocampus, as well as reduced spongiform change. **D**. NeuN counts of pyramidal neurons in NBH versus scrambled-treated and M2D-treated prion mice. M2D confers neuroprotection close to levels seen in NBH mice. **E**. Semi-quantitative scoring of spongiosis in NBH, scrambled-treated and M2D-treated mice. Sections showing no signs of spongiosis were scored 0, severe spongiosis was scored 4, as described (White *et al*, 2008). M2D significantly reduced spongiosis compared to scrambled-treated prion mice, which is absent in uninfected NBH mice. **F**. Total PrP and proteinase K-resistant PrP^Sc^ levels in NBH-injected mice and in prion-diseased mice injected with scrambled- or M2D ASOs. PrP^Sc^ levels are unaffected by M2D-mediated RBM3 induction.

## Materials and Methods

### RNA-Seq analysis and bioinformatics

RNA sequencing data from mouse primary hepatocytes are deposited under GSE158882 (Neumann *et al*., 2020). Sequencing data from HeLa cells after knockdown and rescue of *UPF1, SMG6* and *SMG7* were obtained from SRP083135 (Colombo *et al*., 2017). Mapping of reads to reference genomes (mm10 for mouse, hg38 for human) was performed using STAR version 2.5.3a (Dobin *et al*, 2012). The RBM3 Sashimi plot was generated using a customized version of ggsashimi (Garrido-Martin *et al*, 2018), which additionally displays conservation scores. Percent spliced in values for *Rbm3* E3a after knockdown and depletion was manually calculated from junction read counts.

### Tissue culture cells

HEK293T, HeLa and N2a cell stocks are maintained in liquid nitrogen and early passage aliquots are thawed periodically. To rule out mycoplasma contamination, cell morphology is routinely assessed and monthly checked using a PCR-based assay. HEK293T and HeLa cells were cultured in DMEM high glucose with 10% FBS and 1% Penicillin/ Streptomycin. N2a cells were cultured in 50% OptiMEM / 50% OptiMEM Glutamax with 10% FBS and 1% Penicillin/Streptomycin. Mouse hippocampal neurons were isolated and maintained as previously described (Peretti *et al*., 2021). All cell lines were usually maintained at 37°C and 5% 5% CO_2_. For temperature experiments cells were shifted into pre-equilibrated incubators for the indicated times. For square-wave temperature cycles we used two incubators set to 34°C and 38°C and shifted the cells every 12 hours (Preussner *et al*., 2017). Transfections of HEK293T or N2a cells (for minigenes or CRISPR guides) using Rotifect (Roth) were performed according to the manufacturer’s instructions. Cycloheximide (Sigma) was used at 40μg/ml final concentration or DMSO as solvent control. For morpholino experiments, cells were seeded and transfected 1 day later using Endoporter following the manufacturer’s manual. Morpholinos (*RBM3* 5’ss: GTCTCCCCTGCTACTACTTACATCT and standard control) and Endoporter transfection reagent were purchased from Gene Tools. MOE antisense oligonucleotides (1ul, 100ng/ul) were purchased from Mycrosynth (see Table S2 for sequences) and transfected into Hela or N2a cells (3*10^6^ cells per well on a 12-well plate) via Rotifect, and 4 hours later the transfected cells were transferred to incubators with the indicated temperature, followed by RNA extraction 24 hrs later and RT-qPCR or splicing sensitive PCR.

### RT–PCR and RT–qPCR

RT–PCRs were done as previously described (Preussner *et al*, 2014). Shortly, RNA was extracted using RNATri (Bio&Sell) and 1μg RNA was used in a gene-specific RT reaction. Endogenous *RBM3* splicing was analyzed with a radioactively labeled forward primer in exon 3 (5’-TCATCACCTTCACCAACCCA) and a reverse primer in exon 5 (5’-TCTAGAGTAGCTGCGACCAC). For analysis of minigene splicing, the RNA was additionally digested with DNase I and re-purified. Minigene splicing was investigated with minigene specific primers: T7fwd: 5’-GACTCACTATAGGGAGACCC; BGHrev: 5’-TAGAAGGCACAGTCGAGG. In some cases a minigene specific RT with BGHrev was followed by a PCR with forward primer targeting exon 3 (5’-TGGTTGTTGTCAAGGACCGGG) and reverse primer targeting exon 5 (5’-CTCTAGAGTAACTGCGACCAC). For RT–qPCR, the *RBM3* gene-specific primer was combined with a housekeeping gene reverse primer in one RT reaction. qPCR was then performed in a 96-well format using the Blue S’Green qPCR Kit Separate ROX (Biozym) on Stratagene Mx3000P instruments. qPCRs were performed in technical duplicates, mean values were used to normalize expression to a housekeeping gene (human: GAPDH; mouse: HPRT); DCT, and D(DCT)s were calculated for different conditions.

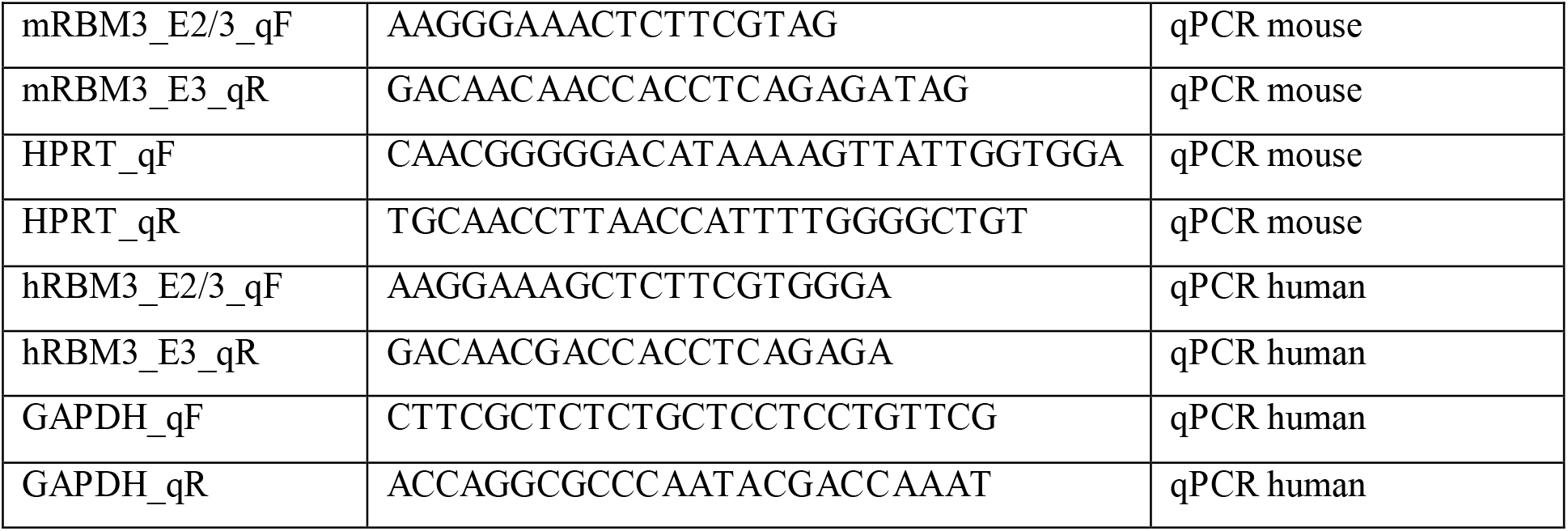

### Generation of CRISPR-Cas9 edited cells and analysis of RBM3 expression

For genome-engineering in HEK293 cells, sequences flanking E3a of *RBM3* were analyzed for sgRNA candidates in silico using the Benchling tool. A pair of oligonucleotides for the highest ranked candidate sgRNA (Ran *et al*, 2013) upstream and downstream of the exon was synthesized and subcloned into the PX459 vector. sgRNA sequences #1: 5’-TGTGTCTGCTCGGGGCAGCG; #2: 5’-CCTGTGAGTGGGCACTGCG; #3: 5’-TCCTGATGAAGCCATTCTG. Cells were co-transfected with guide RNA #3 and either #1 or #2 in 6-well plates using Rotifect (Roth) according to the manufacturer’s instructions. 48 hours after transfection, the transfected cells were selected with 1 ug/ml puromycin and clonal cell lines were isolated by dilution. Genomic DNA was extracted using DNA extraction buffer (200 mM Tris/HCl pH 8.3, 500 mM KCl, 5 mM MgCl2, 0.1% gelatin in H_2_O) and to confirm the exon knockout on DNA level, a genotyping PCR was performed using primers binding in introns upstream and downstream of the cutting sites (FWD: 5’-ATCTGCAGAGGGACCTTGTC; REV: 5’-CAGACTTGCCTGCATGATCC). In promising clones the exon knockout was additionally confirmed after RNA isolation by splicing sensitive PCR using one forward primer in exon 3 and one reverse primer in exon 4. RBM3 total expression levels were investigated by RT-qPCR and Western blot.

### Western blot

Whole-cell extracts (WCEs) were prepared with lysis buffer (20mM Tris (pH 8.0), 2% NP-40 (v/v), 0,01% sodium deoxycholate (w/v), 4mM EDTA and 200mM NaCl) supplemented with protease inhibitor mix (Aprotinin, Leupeptin, Vanadat and PMSF). Concentrations were determined using RotiNanoquant (Roth), according to the manufacturer’s instructions. SDS-PAGE and Western blotting followed standard procedures. Western blots were quantified using the ImageQuant TL software. The following antibodies were used for Western blotting: hnRNPL (4D11, Santa Cruz), RBM3 (proteintech), GAPDH (GT239, GeneTex), actin (4970, Cell Signaling).

### Minigene constructs

Cloning was performed using PCR introducing HindIII and XhoI sites, and ligation into pcDNA3.1 (+). Constructs were cloned using a forward primer in the intron upstream of exon 3 (mouse: 5’-AACTTAAGCTTCTGTGGCTGTGCCTGGCT; human: 5’-AACTTAAGCTTTCCGGCCACCCTTTGCTAC), a reverse primer in the intron downstream of exon 4 (mouse: 5’-CTAGACTCCAGTTCAGACATAGGCTCTTAA; human: 5’-TAGACTCGAGATAGGCAACTCTCCC TCTCA) and human or mouse DNA as template. For each of mRRM3 mutant (see Table S1) and sub-mutant cloning, the mutated sequences were deleted or replaced by sequences from human beta-globin (M3 contains a 3’ss, M6 a 5’ss and the remaining mutants exon 2 sequences). Briefly, two DNA fragments were amplified from the mRBM3 minigene with PCR primer pairs introducing the mutation. Then, these two DNA fragments were used as templates for a PCR to get the full-length mRBM3 mutants. All minigene sequences were confirmed by sequencing (Microsynth SeqLab).

### Primary neuronal culture

Primary neurons were isolated from the hippocampi of both male and female C57Bl6/N mouse pups at post-natal day 0 or 1. For the isolation and culture of hippocampal neurons we followed the protocol of Beaudoin et. al. with slight modifications (Beaudoin *et al*, 2012). Briefly, hippocampi were extracted into Hibernate A media (Gibco) and incubated at 37°C with papain solution for 20 minutes. Papain solution was removed and trypsin inhibitor was added for 5 minutes. Hippocampi were then washed 3 times in pre-warmed plating media (Neurobasal A, B27 supplement, Glutamax, Horse serum, 1M HEPES pH 7.5) before being triturated 8-10 times. The suspension was strained and 800,000 cells were seeded onto 6-well plates coated with poly-L-lysine. Media was changed to neuron media (Neurobasal A, B27 supplement, Glutamax, Penicillin/Streptomycin) 4 hours post seeding. Primary neurons were maintained at 37°C, 5% CO2 or as indicated at 32°C. 5-fluoro-2’-deoxyuridine (Sigma-Aldrich) was added at a final concentration of 7.15 μg/ml to inhibit glial growth (Vossel *et al*, 2015). A third of the media was changed for fresh media every 4-5 days. Experimental procedures lasting 24-48 hours were started at day 19-20 D.I.V. to finish at 21 D.I.V.

### Mice

All animal work conformed to UK Home Office regulations and were performed under the Animal [Scientific Procedures] Act 1986, Amendment Regulations 2012 and following institutional guidelines for the care and use of animals for research. All studies were ethically reviewed by the University of Cambridge Animal Welfare and Ethical Review Body (AWERB). Mice were housed in groups of 2-5 animals/cage, under 12hours light/dark cycle and were tested in the light phase. Water and standard mouse chow were given ad-libitum. Mice were randomly assigned treatment groups by cage number. Experimenters were blind to group allocation during the experiments and when assessing clinical signs. Procedures were fully compliant with Animal Research: Reporting of In Vivo Experiments (ARRIVE) guidelines.

### Prion infection of mice

3-week-old tg37^+/−^ mice were inoculated intra-cerebrally into the right parietal lobe with 30 μL 1% brain homogenate of Chandler/RML (Rocky Mountain Laboratories) prions under general anesthetic, as described (Mallucci *et al*., 2002). Animals were culled when they developed clinical signs of scrapie as defined in (Mallucci *et al*, 2003; Mallucci *et al*., 2002; Mallucci *et al*, 2007). Control mice received 1% normal brain homogenate.

### Administration of ASOs

Uninoculated tg37 hemizygous mice were injected with 100 or 300 μg of scrambled ASO or M2D ASO. ASOs were administered via intracerebroventricular injection at −0.3Y, 1.0X, 3.0Z anterior to bregma. Mice were culled 3 weeks post injection. Prion-infected mice were injected with 200μg of scrambled ASO or M2D ASO at 3 w.p.i and culled at 12 w.p.i. This dose was chosen as an average of the efficacious doses of 100 and 300ug. All surgeries were performed under general anesthesia.

### Histology

Paraffin-embedded brains were sectioned at 5 mm and stained with H&E as described (Moreno *et al*, 2012). Neuronal counts were determined by quantifying NeuN-positive pyramidal CA1 neurons as described (Moreno *et al*., 2012).

### Statistical analysis

Data is presented as the mean ± standard error of the mean (SEM) unless otherwise specified in the legend. Statistical significance was determined using Graphpad Prism v7 or v8, using a Student’s t test or one-way ANOVA with Tukey’s test for multiple comparisons for normally distributed data or Kruskal-Wallis test and Dunn’s multiple comparisons test for non-normally distributed data sets. Statistical significance was accepted at P=<0.05. In the figure legends, “ns” denotes P≥0.05, * denotes P=<0.05, ** denotes P=<0.01, ***P=<0.001 and **** denotes P=<0.0001.

## Acknowledgements

The authors would like to thank HPC Service of ZEDAT, Freie Universität Berlin, for computing time. We thank members of the Heyd lab for constructive discussions and comments.

## Authors contributions

MP performed the bioinformatics analysis with help from AN. AKE generated and analyzed CRISPR/Cas9-mediated knockout clones. DP performed analyses in primary neurons. MZ and MP performed minigene experiments. TH, SS and MZ did the ASO screen. HLS, DH and DS performed in vivo experiments and analyses of ASO effects. FH, MP and GRM designed the study and supervised the work, with initial input from DP. All authors were involved in data interpretation. The manuscript was written by MP, FH, GRM and HLS, with input from all authors.

## Supplementary Materials

Figs. S1 to S4

Tables S1 to S2

