## Supplementary Figure S1-S4 for "ASO targeting temperature-controlled *RBM3* poison exon splicing prevents neurodegeneration in vivo"

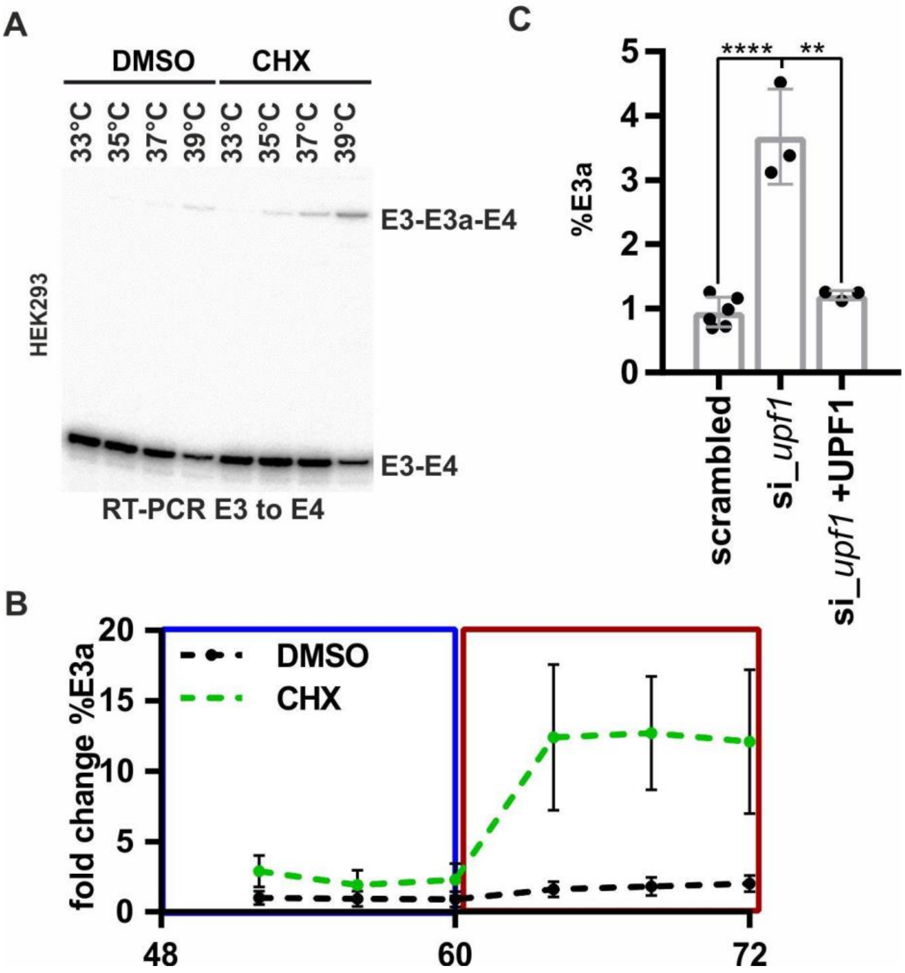


**Figure S1:** ***RBM3*** **intron 3 contains an evolutionary conserved heat-induced poison exon. A.** Representative gel image showing heat-induced and CHX-stabilized formation of the E3a isoform in human HEK293 cells (quantification in Figure 1C). **B.** Rhythmic *RBM3* E3a regulation. HEK293 cells were pre-entrained with square-wave temperature cycles (12h 34°C/ 12h 38°C) for 48h. For the last 24h, cells were treated with DMSO or CHX every 4h and harvested after 4h and analyzed by splicing sensitive RT-PCR (n=6, mean ± SD). **C.** *RBM3* E3a stabilization in response to *UPF1* knockdown and rescue (mean +/- SD, n=3, all individual data points are shown). Sequencing data were obtained from SRP083135[26]. Statistical significance was determined by unpaired t-test and is indicated by asterisks: p values: **p<0.01, ****p<0.0001. Data from (*29*).

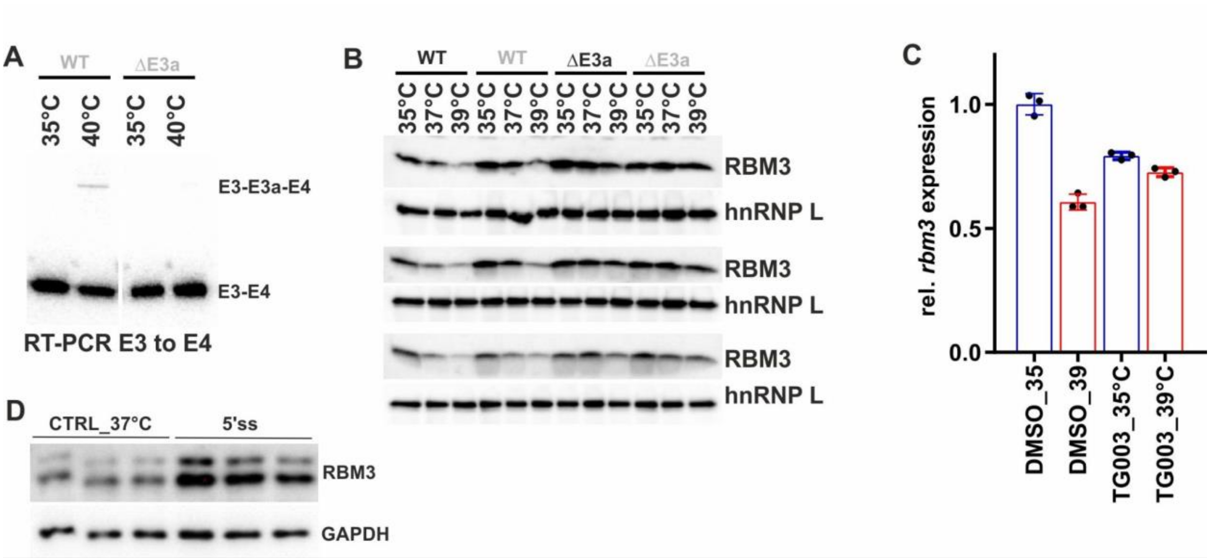


**Figure S2: E3a controls temperature dependent RBM3 expression. A.** An RT-PCR in CHX-treated HEK293 cells confirms CRISPR/CAS9-mediated removal of E3a in a DE3a clone at the RNA level. **B.** Western blot analysis of RBM3 protein expression, hnRNP L served as a loading control. Two independent WT and DE3a clones are shown in biological triplicates. Quantification in Figure 2C. **C.** Inhibition of CLK1/4 kinase by TG003 abolishes the effect of temperature on *rbm3* expression. Whippet derived TpM values are shown relative to DMSO 35°C. This reveals an almost 2-fold difference in *rbm3* levels comparing 6h DMSO 35°C vs 39°C. Note that this is basically abolished by adding TG003 during the shift from 39°C to 35°C. Data from (*8*). **D.** Blocking *rbm3* E3a inclusion induces RBM3 protein levels. Primary hippocampal neurons were transfected with the indicated MOs for 48 hours and investigated by Western blotting. GAPDH served as a loading control. Quantification in Figure 2E.

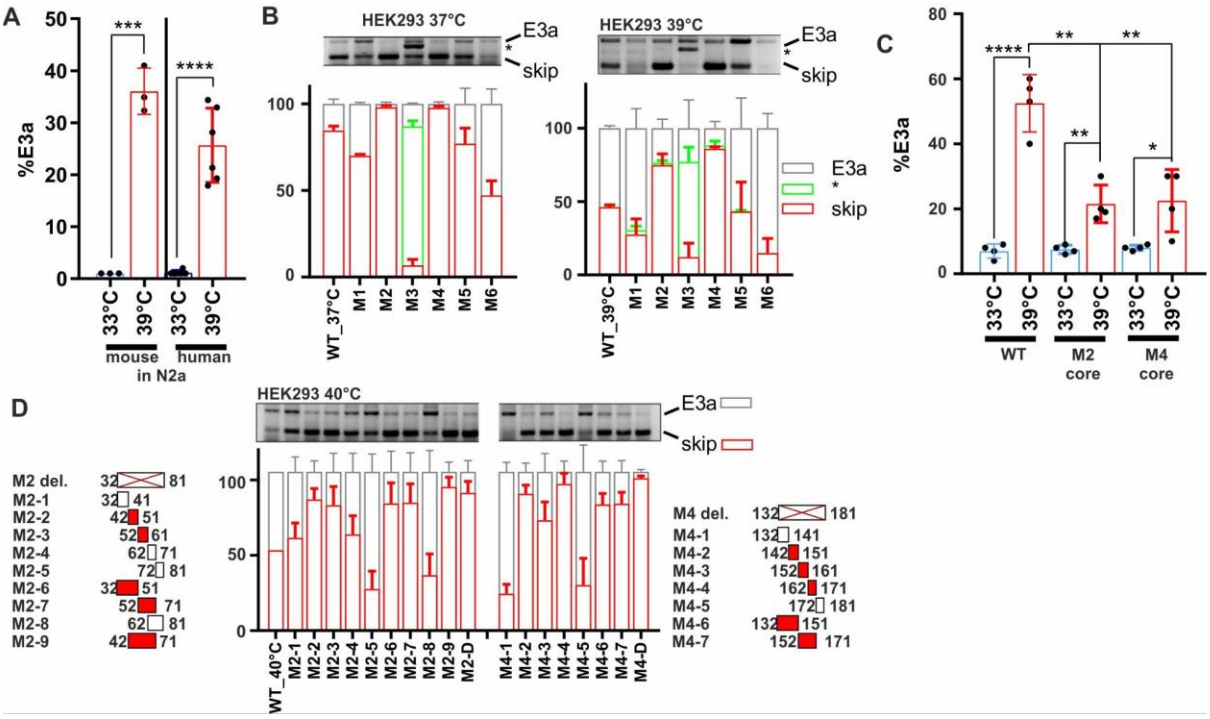


**Figure S3: Mapping *cis*-regulatory elements controlling *RBM3* E3a inclusion. A.** An *RBM3* minigene reproduces temperature controlled E3a inclusion. Human and mouse minigenes were transfected into N2a cells and analysed as in Figure 3A. **B.** Systematic mutational screening for regulatory elements (see Table S1). The indicated sequences were replaced by sequences from human beta-globin. M2, M4 and M5 contain the same exon 2 sequence from beta-globin exon 2. Only in the M2 and M4 context these sequences prevent inclusion, ruling out the possibility that we included a silencer element. M1 and M3 contain sequences of the beta-globin 3'ss, M6 the beta-globin 5'ss (these mutations do not result in an increased splice site strength). E3a inclusion in HEK293 at 37°C and 39°C was investigated by minigene specific RT-PCR. On top a representative gel is shown. Below, quantifications of the detected isoforms (n=2). Note that replacing the internal 3'ss with a globin 3'ss promotes its usage. **C.** Temperature response of the indicated minigenes. We deleted the evolutionary conserved core of the M2 and M4 regions. Briefly, M2-2 and M2-3 are 100% conserved between human and mouse. Thus, M2-2 and M2-3 are regarded as core sequence of the M2 enhancer to be deleted in hRBM3 minigene. For the M4 region, M4-3 is the central region of the conserved sequence. Therefore, M4-3 and a part of the upstream sequence of M4-4 were deleted as core sequence of M4 mutant in the hRBM3 minigene (mean ± SD, n=4). **D.** Detailed mutational screening of the M2 region (borders indicated on the left) and M4 region (borders indicated on the right). See Table S1 for sequences. In M2 or M4 del. the M2 or M4 sequences are removed (and not replaced). In M2-1 to M2-9 and M4-1 to M4-7 the indicated sequences are replaced by human beta-globin exon 2 sequences from the same relative position. On top a representative PCR image is shown. Below, quantifications of the detected isoforms (n=2).

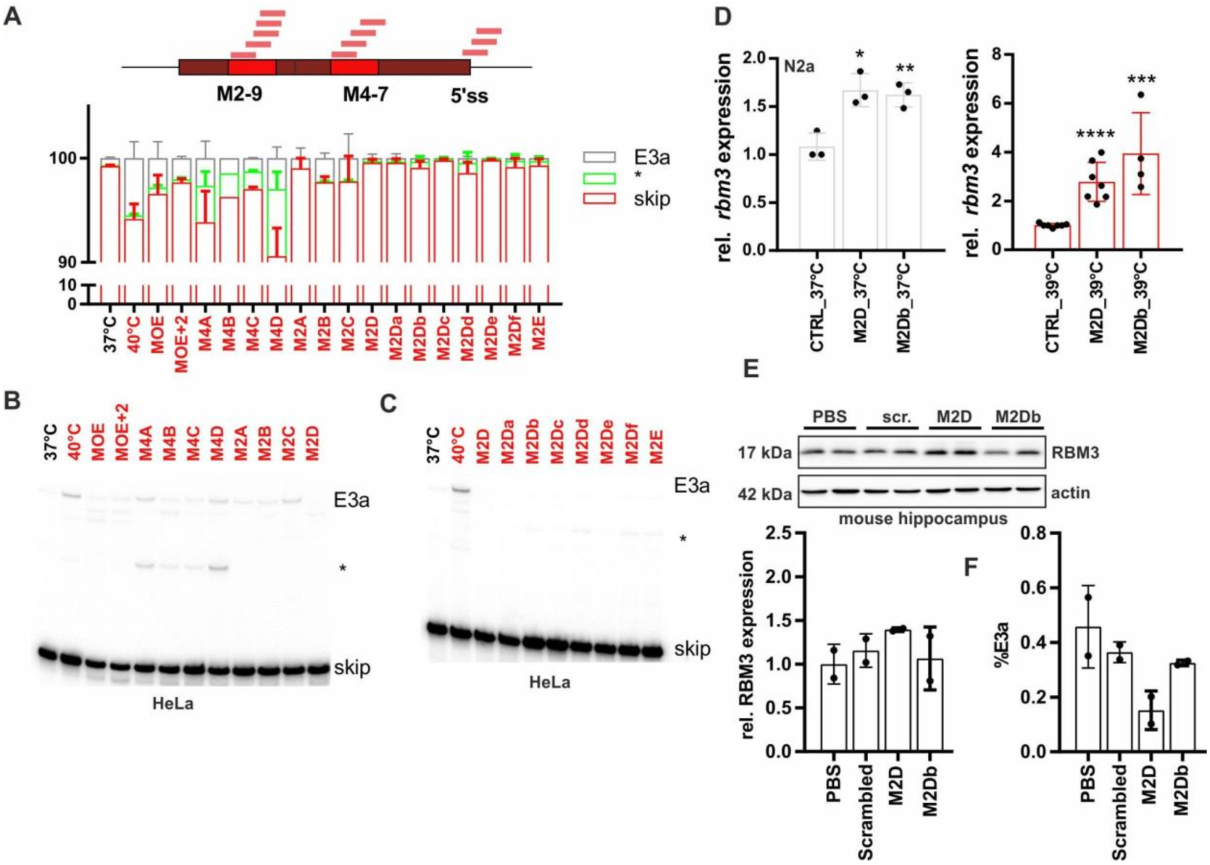


**Figure S4: Screening for ASO sequences that mediate *RBM3* E3a exclusion. A-C.** ASOs targeting M2-9, M4-7 or the 5'ss (see Table S2) prevent *RBM3* E3a inclusion in human HeLa cells. ASOs were transfected and cells were kept at 40°C for 24 hours. Control samples at 37°C and 40°C are shown. All samples were treated with CHX for the last 4 hours. In B and C representative gels are shown. Exon 3a inclusion was investigated by splicing sensitive radioactive RT-PCR. A quantification is shown in A (n=2; n=5 for all M2D variants and M2E, n=1 for M4B). The asterisk marks the use of internal 5' and 3'ss that is promoted by all ASOs targeting the M4 region. Note that all variants targeting the M2D region almost quantitatively prevent E3a inclusion and, at 40°C, lead to inclusion levels that are lower than the one observed at 37°C for control cells. **D.** M2D and M2Db induce *rbm3* mRNA expression in mouse N2a cells. ASOs were transfected for 24 hours at 37°C (left) or at 39°C (right). *rbm3* induction was measured relative to a CTRL ASO and relative to HPRT expression (mean ± SD, n>=3, all individual data points are shown; unpaired t-test derived p-value *p<0.5, **p<0.01, ***p<0.001, ****p<0.0001). **E.** M2D, but not M2Db, induces RBM3 protein expression *in vivo* (100µg dose per mouse). Hippocampus samples from two independent mice per condition were analyzed by Western blotting (top) and RBM3 signal was quantified relative to actin and PBS (bottom, n=2). **F.** M2D, but not M2Db, reduces E3a inclusion *in vivo* (300µg dose per mouse). Cerebellum RNA samples from two independent mice per condition were analyzed by splicing sensitive RT-PCR and %E3a signal was quantified (n=2).
